# Structure and mechanistic features of the prokaryotic minimal RNase P

**DOI:** 10.1101/2021.05.07.443126

**Authors:** Rebecca Feyh, Nadine B. Wäber, Simone Prinz, Pietro Ivan Giammarinaro, Gert Bange, Georg Hochberg, Roland K. Hartmann, Florian Altegoer

## Abstract

Endonucleolytic removal of 5’-leader sequences from tRNA precursor transcripts (pre-tRNAs) by RNase P is essential for protein synthesis. Beyond RNA-based RNase P enzymes, protein-only versions of the enzyme exert this function in various Eukarya (there termed PRORPs) and in some bacteria (*Aquifex aeolicus* and close relatives); both enzyme types belong to distinct subgroups of the PIN domain metallonuclease superfamily. Homologs of *Aquifex* RNase P (HARPs) are also expressed in some other bacteria and many archaea, where they coexist with RNA-based RNase P and do not represent the main RNase P activity. Here we solved the structure of the bacterial HARP from *Halorhodospira halophila* by cryo-EM revealing a novel screw-like dodecameric assembly. Biochemical experiments demonstrate that oligomerization is required for RNase P activity of HARPs. We propose that the tRNA substrate binds to an extended spike-helix (SH) domain that protrudes from the screw-like assembly to position the 5’-end in close proximity to the active site of the neighboring dimer subunit. The structure suggests that eukaryotic PRORPs and prokaryotic HARPs recognize the same structural elements of pre-tRNAs (tRNA elbow region and cleavage site). Our analysis thus delivers the structural and mechanistic basis for pre-tRNA processing by the prokaryotic HARP system.

## Introduction

Ribonuclease P (RNase P) is the essential endonuclease that catalyzes the 5’-end maturation of tRNAs (Klemm et al. 2016; Schencking, Rossmanith, and Hartmann 2020; Guerrier-Takada et al. 1983). The enzymatic activity is present in all forms of life yet shows a remarkable variation in the molecular architecture. There are two basic types of RNase P, RNA-based and protein-only variants. The former consist of a structurally conserved, catalytic RNA molecule that associates with a varying number of protein cofactors (one in Bacteria, 5 in Archaea, 9-10 in Eukarya (Klemm et al. 2016; Jarrous and Gopalan 2010). Protein-only enzymes arose independently twice in evolution. In Eukarya, a <u>p</u>rotein-<u>o</u>nly <u>R</u>Nase <u>P</u> (termed PRORP) apparently originated at the root of eukaryotic evolution and is present in four of the five eukaryotic supergroups (Lechner et al. 2015). There, this type of enzyme replaced the RNA-based enzyme in one compartment or even in all compartments with protein synthesis machineries, such as land plants harboring PRORP enzymes in the nucleus, mitochondria and chloroplasts (Gobert et al. 2010). In metazoan mitochondria, PRORP requires two additional protein cofactors for efficient function (Holzmann et al. 2008).

More recently, a bacterial protein-only RNase P, associated with a single polypeptide as small as ~23 kDa, was discovered in the hyperthermophilic bacterium *Aquifex aeolicus* that lost the genes for the RNA and protein subunits (*rnpB* and *rnpA*) of the classical and ancient bacterial RNase P (Nickel et al. 2017). This prokaryotic type of minimal RNase P system was named HARP (for: <u>H</u>omolog of <u>A</u>quifex <u>RN</u>ase <u>P</u>) and identified in 5 other of the 36 bacterial phyla beyond Aquificae (Nickel et al. 2017). Among HARP-encoding bacteria, only some lack the genes for RNA-based RNase P (i.e., Aquificae, Nitrospirae), while others harbor *rnpA* and *rnpB* genes as well (Daniels et al. 2019; Nickel et al. 2017). Overall, HARP genes are more abundant in archaea than bacteria. However, all of these HARP-positive archaea also encode the RNA and protein subunits of the RNA-based RNase P (Nickel et al. 2017; Daniels et al. 2019). Remarkably, HARP gene knockouts in two Euryarchaeota, *Haloferax volcanii* and *Methanosarcina mazei*, showed no growth phenotypes under standard conditions, temperature and salt stress (*H. volcanii*) or nitrogen deficiency (*M. mazei*) (Schwarz et al. 2019). In contrast, it was impossible to entirely erase the RNase P RNA gene from the polyploid genome of *H. volcanii* (~18 genome copies per cell in exponential growth phase; (Breuert et al. 2006)). Even a knockdown to ~20% of the wild-type RNase P RNA level in *H. volcanii* was detrimental to tRNA processing and resulted in retarded cell growth (Stachler and Marchfelder 2016). The findings suggest that HARP is neither essential nor represents the housekeeping RNase P function in Archaea, explaining its sporadic loss in archaea. HARPs are evolutionarily linked to toxin-antitoxin systems (Daniels et al. 2019; Schwarz et al. 2019; Gobert, Bruggeman, and Giegé 2019). Frequently, the toxin proteins are endoribonucleases that cleave mRNA, rRNA, tmRNA or tRNA to inhibit protein biosynthesis in response to certain stresses (Masuda and Inouye 2017). Conceivably, the progenitor of *A. aeolicus* and related Aquificaceae might have acquired such a toxin-like tRNA endonuclease via horizontal gene transfer and established it as the main RNase P activity with relatively little reprogramming.

HARPs belong to the PIN domain-like superfamily of metallonucleases. They were assigned to the PIN_5 cluster, VapC structural group, whereas eukaryal PRORPs belong to a different subgroup of this superfamily (Matelska, Steczkiewicz, and Ginalski 2017; Gobert, Bruggeman, and Giegé 2019). HARPs oligomerize and Aq880 was originally observed to elute as a large homo-oligomeric complex of ~420-kDa in gel filtration experiments (Nickel et al. 2017). However, its specific mode of substrate recognition and the underlying structural basis is lacking to date. Here, we present the homo-dodecameric structure of the HARP from the *γ*-bacterium *Halorhodospira halophila* SL1 (Hhal2243) solved by cryo-EM at 3.37 Å resolution. Furthermore, we employed mass photometry to investigate the oligomerization behavior of HARP and correlated the oligomeric state with enzyme activity. Our structure reveals that HARPs form stable dimers via a two-helix domain inserted into the metallonuclease domain. These dimers further assemble into a screw-like assembly resulting in an asymmetric and thus imperfect novel type of homo-dodecamer. Our biochemical analysis suggests that pre-tRNA processing involves the neighboring dimer subunits and thus requires the presence of a higher HARP oligomer. In conclusion, we here present the structural basis for the RNase P-like pre-tRNA processing activity of prokaryotic HARPs.

## Results

### Dodecameric structure of the HARP from *Halorhodospira halophila*

Structural information on *A. aeolicus* RNase P (Aq880) and HARPs is lacking and a mechanistic understanding of pre-tRNA processing by HARPs has remained unknown so far. As previously observed by size exclusion chromatography (SEC), Aq880 forms large oligomers of ~420 kDa (Nickel et al. 2017). Our attempts to resolve its structure by X-ray crystallography or NMR were unsuccessful. We also purified other HARPs to increase the chances of successful structure determination. Among those was the HARP of *Halorhodospira halophila* (Hhal2243) that was purified to homogeneity using a two-step protocol consisting of Ni^2+^-affinity chromatography and anion-exchange chromatography (**Fig. S1**; see Materials and Methods). Hhal2243 formed an assembly of similar size as Aq880 (**Fig. S2A**) but adopted a more uniform oligomeric state than Aq880 (see below). Like Aq880 (Marszalkowski, Willkomm, and Hartmann 2008), Hhal2243 showed pre-tRNA processing activity in the presence of Mg^2+^ and Mn^2+^ (**Fig. S2B, C**).

We succeeded in solving the structure of Hhal2243 by cryo-electron microscopy (cryo-EM) to 3.37 Å resolution (**Figs. 1, S3, Tab. S1**). Hhal2243 assembles, in a left-handed screw-like manner, into a dodecamer with each molecule being rotated by approximately 58°. The first and last subunits of one screw turn are separated by a slightly larger angle of 70° (**Fig. 1A**). Hhal2243 consists of a PIN-like metallonuclease domain into which two helices are inserted that we termed the “spike helix” (SH) domain (**Fig. 1B**). The PIN-like domain is formed by six α-helices (α1-α4, α7, α8) and four β-strands (β1-β4) that fold into a α/β/α domain with a central, four-stranded parallel β-sheet (**Fig. 1C**). Two Hhal2243 monomers align head-to-head with their SH domains consisting of helices α5 and α6 to form a dimer, while two SH domains form a four-helix bundle resulting in six spikes that protrude from the dodecameric assembly (**Fig. 1C**). The dimer interface covers a buried surface area of 1300 Å^2^ and is mainly of hydrophobic nature with two clamping salt bridges formed by R141 and E91 from either monomer, respectively (**Fig. 1D, E**).

**Figure 1.**
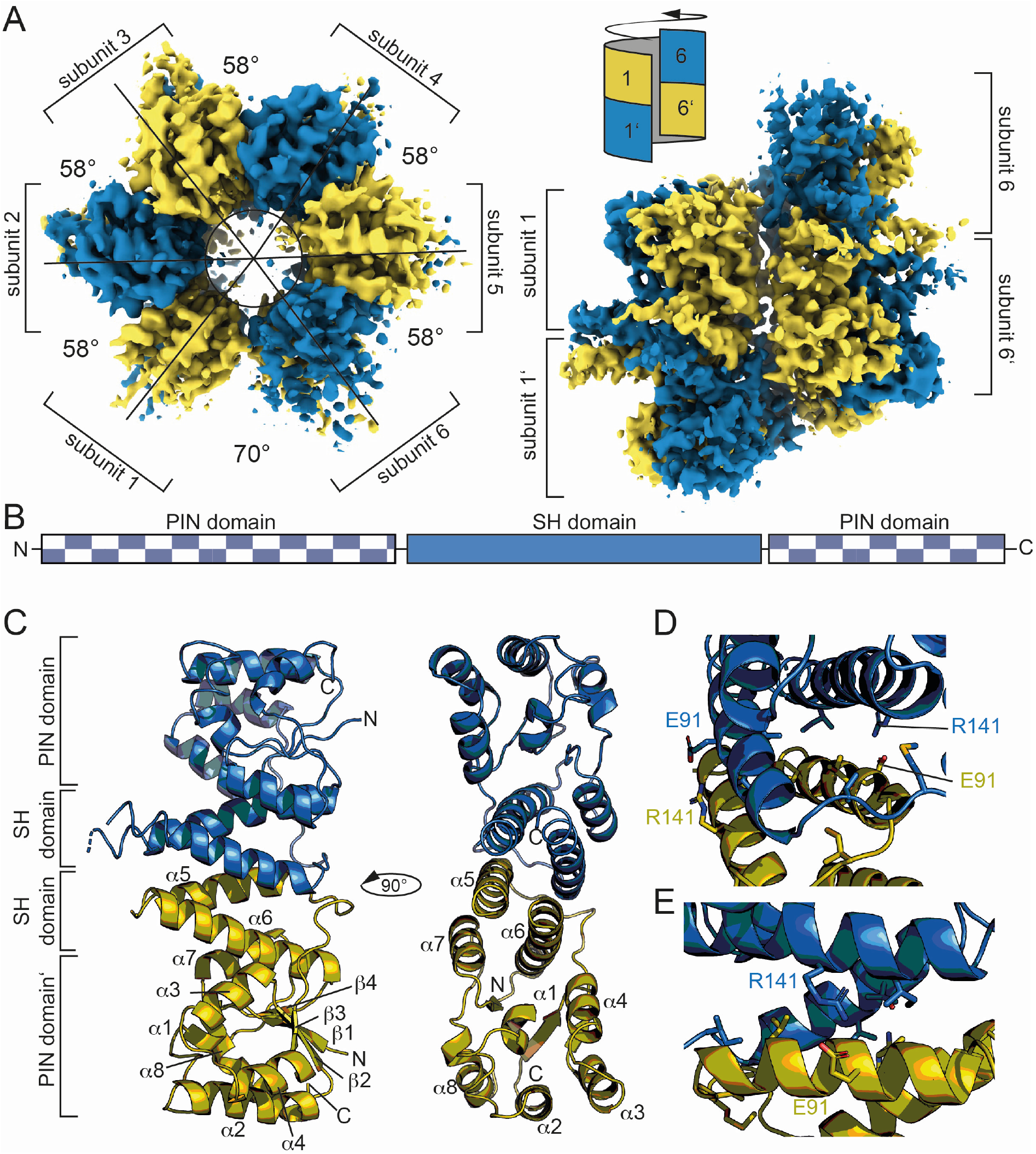
Dodecameric structure of Hhal2243. **A.** Cryo-EM electron density map of Hhal2243 shown from a top (left) and a side view (right). The monomer subunits are colored in blue and olive, respectively. The numbers indicate the angles between the dimer subunits. The sketch in the upper left corner of the view on the right indicates how the subunits assemble to form the screw-like arrangement of the dodecamer. **B.** Domain architecture of Hhal2243. **C.** Model of the Hhal2243 dimer. The protein consists of a PIN 5 domain with an inserted spike-helix (SH) domain forming the dimer interface. **D.** and **E.** Detailed view of the dimer interface. The clamping salt bridges are shown as sticks.

### Oligomerization and its influence on pre-tRNA processing activity

Our findings and the conservation of HARPs suggested that the dodecameric superstructure represents a conserved feature of HARPs and might therefore be required for their RNase P-like activity. To scrutinize this idea, we took a closer look at the interactions between two dimers. The covered interface extends over 900 Å^2^ and involves the long α4-α5 loop of one subunit and the α7-α8 loop as well as helix α8 of the respective other subunit (**Fig. 2A**). The interactions between interdimer residues are mainly of polar nature.

**Figure 2.**
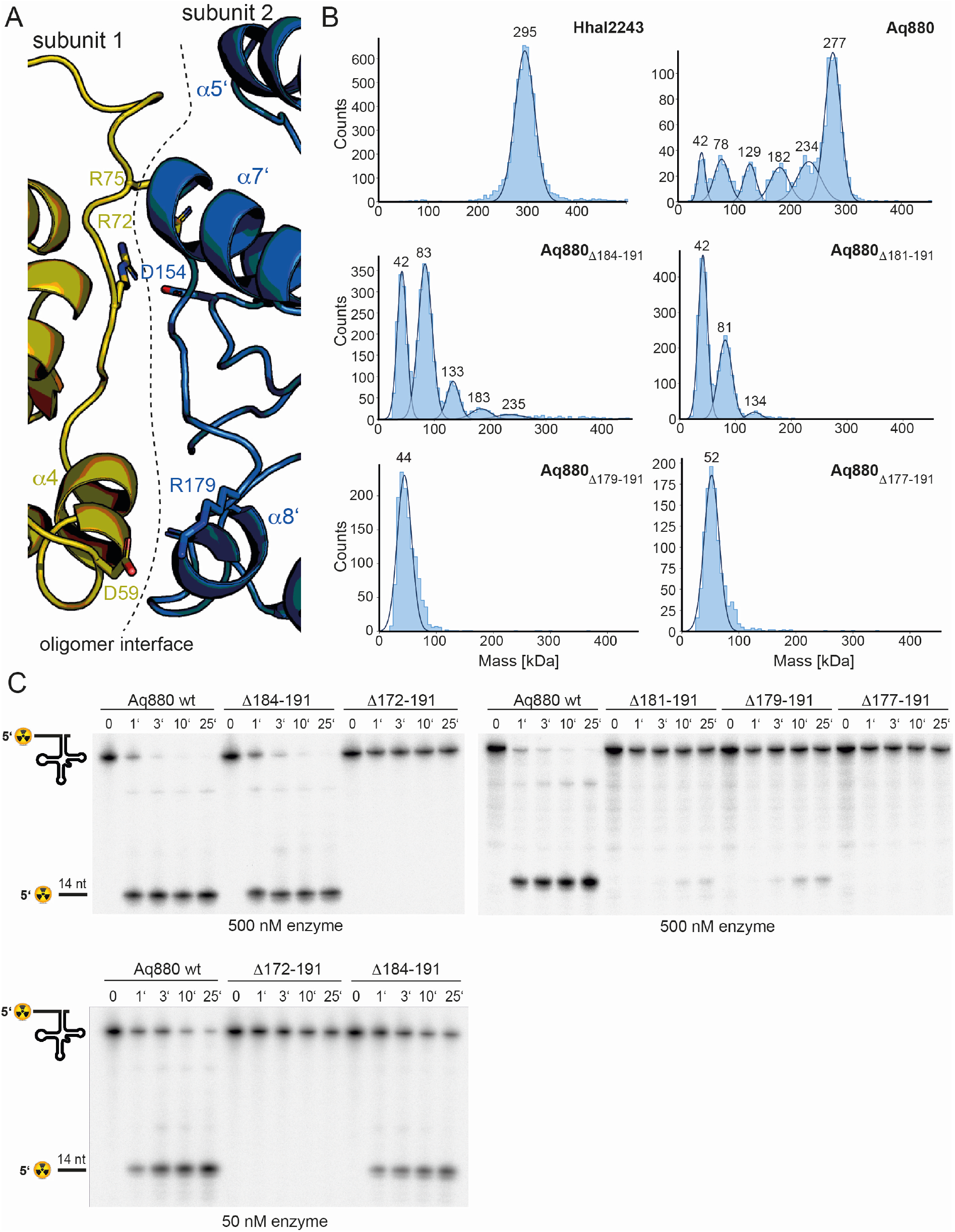
Oligomerization is required for HARP activity. **A.** Oligomer interface between two subunits colored in blue and olive, respectively. **B.** Mass photometry of Hhal2243, Aq880 wt, Aq880_Δ184-191_, Aq880_Δ181-191_, Aq880_Δ179-191_ and Aq880_Δ177-191_. Molecular masses corresponding to the respective gaussian fits are shown in kDa above the fits. **C.** Processing of pre-tRNA^Gly^ by Aq880 wt and derived mutant variants. Aliquots were withdrawn at different time points (1, 3, 10 or 25 min) of incubation at 37 °C; 0, substrate without addition of enzyme. Aq880 wt in comparison with the C-terminal deletion variants Δ172-191, Δ177-191, Δ179-191, Δ181-191 and Δ184-191, all at 500 nM enzyme. For more details, see Materials and Methods.

To investigate the oligomerization behavior we employed mass photometry, a method allowing for the rapid and reliable determination of the dynamic oligomeric distribution of macromolecules in solution (Sonn-Segev et al. 2020; Soltermann et al. 2020). Application of this method to Hhal2243 revealed a stable dodecameric assembly of 295 kDa that included 98 % of all molecules in the sample (**Fig. 2B**, top left panel). Interestingly, despite their similar behavior on SEC, mass photometry of Aq880 unveiled a major species at 277 kDa that, however, included only 47 % of all molecules, while several subspecies with lower molecular weights became visible (**Fig. 2B**, top right panel; **Tab. S3**). This polydispersity of Aq880, not detectable in SEC profiles, is probably the reason why all structural approaches failed so far in the case of Aq880.

To investigate whether oligomerization of Aq880 might impact its pre-tRNA processing activity, we analyzed the oligomer interface of the Hhal2243 structure in more detail. As outlined above, the α4-α5-loop of one subunit is the central element of the interdimer interaction and truncation will most likely impact protein stability (**Fig. 2A**). Thus, we generated several truncations in the C-terminal α8 helix of Aq880, which represents the interacting region on the respective other subunit (**Fig. 2A**). We evaluated the ability of the truncated proteins to oligomerize by mass photometry and their activity in pre-tRNA processing experiments. Five truncated variants of Aq880 were purified to homogeneity and analyzed by SEC (**Fig. S4**). Notably, SEC calibration suggested molecular masses of 55 to 68 kDa for all truncated Aq880 variants (**Fig. S4**). Mass photometry of protein variant Aq880_Δ184-191_ still showed a considerable subfraction of higher-order oligomers (approx. 18 %; compare **Tab. S3**), whereas Aq880_Δ181-191_ formed only 4% hexamer (peak at 134 kDa), 41% tetramer (peak at 81 kDa) beyond the dimer subfraction at 42 kDa (**Fig. 2B**, middle panels; **Tab. S3**). Further truncation of Aq880 had an impact on protein stability, as judged by lower purification yields, and mass photometry revealed that variants Aq880_Δ179-191_ and Aq880_Δ177-191_ form only dimeric species (**Fig. 2B**, lower panels). Processing assays revealed that deletion of residues 184-191 resulted in a protein that still had substantial activity, while all the other truncations showed almost no activity or were entirely inactive (**Table 1**, **Fig. 2C**). Thus, our experiments show that the activity of the enzyme in pre-tRNA processing assays depends on its ability to form higher-order oligomers.

**Table 1:** Processing activities of Aq880 variants

| Aq880 variant | $k_{\text{obs}}$ (min<sup>-1</sup>)<br>(± SD) | substrate cleaved<br>after 25 min (in %) |
| --- | --- | --- |
| wt 50 nM | 0.62 ± 0.15 | 95 |
| wt 500 nM | 2.06 ± 0.42 | 98 |
| Δ184-191 50 nM | 0.22 ± 0.08 | 46 |
| Δ184-191 500 nM | 1.41 ± 0.07 | 97 |
| Δ181-191 500 nM | (4 ± 1) $\times 10^{-3}$ | 9 |
| Δ179-191 500 nM | (7 ± 3) $\times 10^{-3}$ | 15 |
| Δ177-191 500 nM | -- | n.d. |
| Δ172-191 500 nM | -- | n.d. |
| R125A 50 nM | (15 ± 3) $\times 10^{-4}$ | 2 |
| R125A 500 nM | (16.1 ± 0.3) $\times 10^{-2}$ | 26 |
| R129A 500 nM | -- | n.d. |
| R125A/R129A 500 nM | -- | n.d. |
| K119A/R123A/<br>R125A/K127A/R129A 500 nM | -- | n.d. |

n.d., no cleavage detectable

### The active site is conserved among HARPs and PRORPS

Phylogenetic analyses indicate that the two protein-only RNase P systems found in bacteria and eukarya evolved independently (Lechner et al. 2015; Nickel et al. 2017). This is consistent with the different structural basis of the two types of enzymes acting on tRNAs. PRORPs are composed of a PPR domain important for substrate recognition, a central zinc finger domain and a flexible hinge connecting it to the metallonuclease domain (Howard et al. 2012; Teramoto et al. 2020). In contrast, HARPs solely consist of a metallonuclease domain (**Fig. S5A**). The active sites of both molecules superpose well with a root mean square deviation (r.m.s.d) of 0.486 over 30 Ca atoms. In earlier studies we could already show that the catalytic aspartates D7, D138, D142 and D161 are indispensable for Aq880 activity (Nickel et al. 2017; Schwarz et al., 2019). Our Hhal2243 structure supports the prediction that these residues constitute the active site of the protein (**Fig. S5B**), including an almost perfect superposition with three of the four active site aspartates of *Arabidopsis thaliana* PRORP1 (*At*PRORP1; D399, D475 and D493; Howard et al. 2012) (**Fig. S5B**). We conclude that the mechanism of catalysis is conserved among the two distinct protein families with RNase P activity.

### The SH domain is critical for tRNA binding

In a next step, we sought to identify those amino acid residues that are required for the interaction of the enzyme with its pre-tRNA substrate. Helices α5 and α6 of the SH domain expose several conserved arginines and lysines that might be critical for pre-tRNA binding. Notably, several of the residues lie within the distal part of the SH domain that was not resolved in our cryo-EM structure, likely due to a high degree of conformational flexibility. As secondary structure predictions indicated a continuation of the helical arrangement in this region, we generated a homology model of Aq880 based on the Hhal2243 structure and the secondary structure prediction (**Fig. S6A**). The resulting Aq880 model was further verified by rigid body docking into the electron density map of our Hhal2243 structure (**Fig. S6B**).

All residues that were varied to alanines are within the distal part of helix α6 of the SH domain (**Fig. 3A**). Variation of the entire positive arginine/lysine stretch resulted in the inability of the mutant protein to cleave off the 5’-leader sequence (**Fig. 3B**). To narrow down the critical residues, we generated the R125A/R129A double mutant and protein variants with single mutations of R125 or R129. While both R129A and the double mutant R125A/R129A were inactive, R125A retained residual activity (**Fig. 3C**, **Table 1**). Our data suggest that the cluster of positively charged side chains in the SH domain is required for pre-tRNA binding. Although this part is less ordered in our cryo-EM structure it might become more rigid upon tRNA substrate binding.

**Figure 3.**
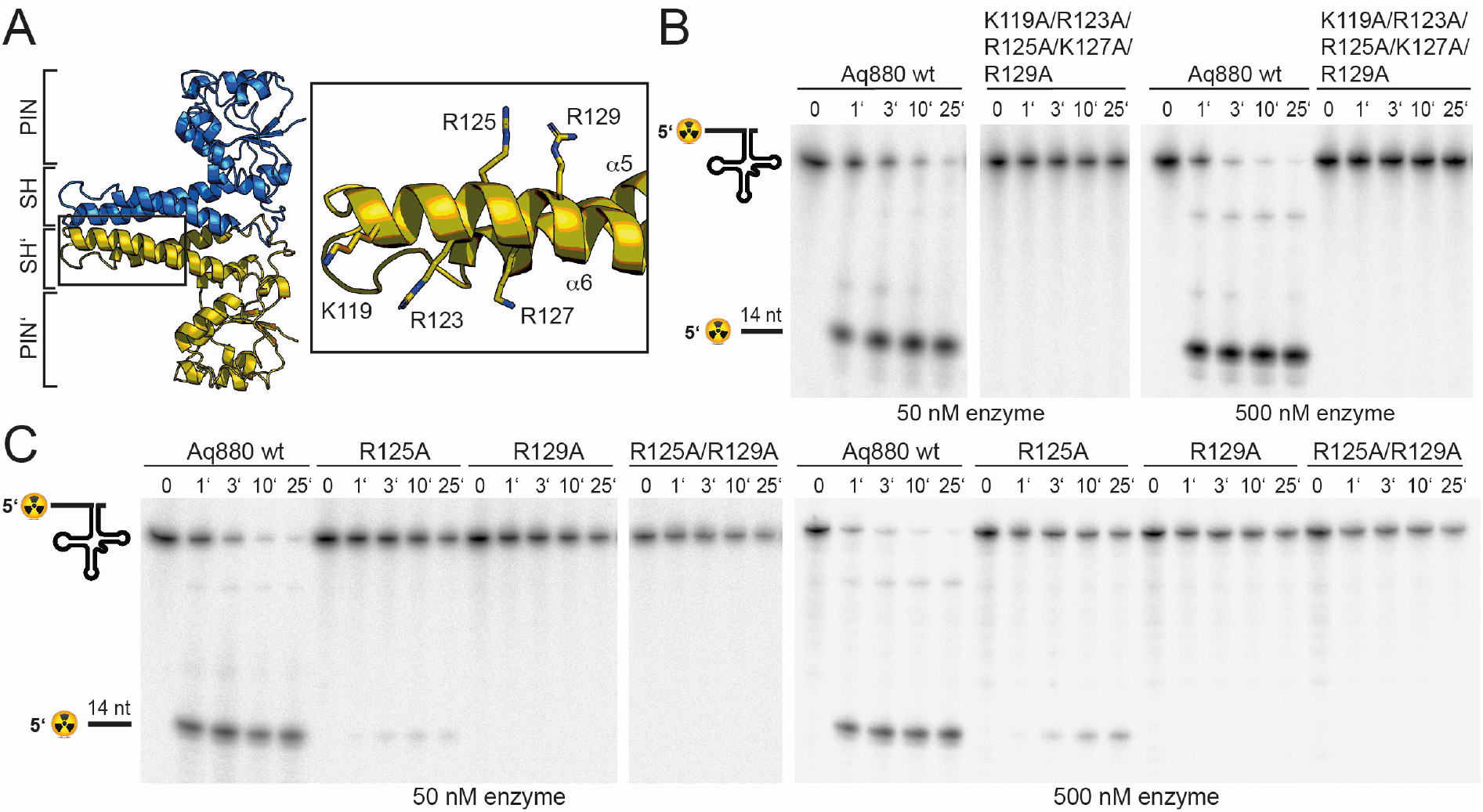
The SH domain is essential for tRNA binding. **A.** Homology model of Aq880 with residues critical for pre-tRNA processing activity in the SH domain displayed as sticks. Processing of pre-tRNA^Gly^ by Aq880 wt and derived mutant variants. Aliquots were withdrawn at different time points (1, 3, 10 or 25 min) of incubation at 37 °C; 0, substrate without addition of enzyme. **B**. Aq880 wt and the quintuple mutant K119A/R123A/R125A/K127A/R129A at 50 nM (left) or 500 nM (right) enzyme; **C**. Aq880 wt, the single mutants R125A and R129A, and the double mutant R125A/R129A, assayed at 50 nM (left) or 500 nM (right) enzyme.

## Discussion

Here, we present the cryo-EM structure of Hhal2243, a member of the recently described HARP family of bacterial and archaeal proteins with RNase P activity (Nickel et al. 2017). Our combined structural and biochemical analysis sheds light on this prokaryotic minimal protein-only RNase P system.

The Hhal2243 HARP structure assembles into a homo-dodecameric ultrastructure. The dodecamer consists of six dimers, composed of monomers aligned head-to-head, which oligomerize in a screw-like fashion (**Fig. 1**). There are many examples of proteins forming symmetric homo-dodecameric assemblies (e.g. Glutamine Synthase (Van Rooyen et al. 2011); Helicases (Bazin et al. 2015) or bacterial DNA-binding proteins expressed in the stationary phase (DPS) (Roy et al. 2008)) and recent studies have discussed the theoretical types of possible quaternary structures (Laniado and Yeates 2020; Ahnert et al. 2015). However, the screw-like assembly of HARPs leads to an asymmetric and thus imperfect novel type of oligomer. This way, HARPs form a defined quaternary structure that is not entirely symmetric, while the screw-like assembly terminates incorporation of new subunits at the stage of a dodecamer. By stepwise truncation of the C-terminus of Aq880, oligomerization could be reduced or abolished, which correlated with functional losses (**Fig. 2C**). Our results thus clearly show that only oligomeric species of HARP are able to cleave the 5’-leader sequence of pre-tRNAs.

HARPs belong to the PIN domain-like superfamily of metallonucleases and share this classification with the eukaryotic PRORPs although the two systems belong to distinct subgroups (Matelska, Steczkiewicz, and Ginalski 2017). We compared our structure to the PIN domain of PRORPs and could demonstrate that the active site residues superpose well (**Fig. S5B**), suggesting that the catalytic mechanism is basically conserved. Notably, the two proteins had to be superposed over a small range of Ca-atoms to obtain a good r.m.s.d. (**Fig. S5A**) owing to the large overall differences of the two PIN domains.

With the knowledge of the structural conservation of the active site, we examined the pre-tRNA substrate recognition and binding by the two protein-only RNase P systems: The crystal structure of the PPR domain of *At*PRORP1 in complex with a tRNA unveiled how PRORPs recognize the pre-tRNA via its PPR RNA-binding domain (Teramoto et al. 2020). The characteristic feature of classical Pentatricopeptide repeats (PPR) is that each PPR repeat recognizes a specific nucleotide, allowing high RNA target specificity (Yan et al. 2019). Interestingly, the PPR domain of PRORP recognizes structural elements of tRNAs rather than single nucleotides in a binding mode reminiscent to that of the RNA-based RNase P holoenzyme (Teramoto et al. 2020). More precisely, the elbow region, where D- and T-loop interact, is the main tRNA-protein interaction interface (Teramoto et al. 2020). The same region, representing the most conserved structural element of tRNAs, is also recognized by RNA-based RNase P enzymes (reviewed in: (Schencking, Rossmanith, and Hartmann 2020)). We thus considered it likely that pre-tRNA recognition by HARPs involves the tRNA elbow region in a similar manner as for all other RNase P types, although we were puzzled by the absence of any evident RNA binding motifs in HARPs. We then focused on exposed positively charged residues and were indeed able to identify a stretch of conserved arginine and lysine residues within the SH domain of the protein. Variation of this positive stretch to alanines rendered the protein inactive (**Fig. 3**).

Combining our knowledge on the active site, on the residues critical for pre-tRNA binding and the correlation of an oligomeric assembly with enzymatic activity, we set out to propose a model for pre-tRNA binding by HARP proteins. Assuming that the outer SH domain, harboring the stretch of positively charged amino acid side chains, interacts with the tRNA elbow, then the distance of approximately 45 Å between the outer SH domain of one dimer and the active site of the neighboring dimer perfectly positions one pre-tRNA molecule for cleavage (**Fig. 4A**). To test this, we used the yeast tRNA from the PRORP structure as model (Teramoto et al. 2020) and docked it onto our HARP structure (**Fig. 4B**). Notably, in this scenario the D-loop is near helix α3 (**Fig. 4B**). We thus also consider it likely that residues within α3 are involved in coordination of the D-loop. Our proposed tRNA binding mode considers that oligomerization is strictly required for HARP activity. This framework enables the cooperation of two adjacent dimer subunits, where one dimer binds the tRNA elbow in such a manner that the pre-tRNA cleavage site, at a distance of ~45 Å, directly reaches into the active site of the neighboring dimer (**Fig. 4B**). This way, five pre-tRNAs can potentially be docked onto one half of the dodecamer (**Fig. 4C**). Although the model is hypothetical at present, other pre-tRNA binding modes seem unlikely based on the spatial constraints imposed by the dodecameric HARP structure and the uniformity and rigidity of prokaryotic tRNAs. Our data furthermore suggest that in the higher oligomeric assemblies the dimers are stabilized by each other for efficient pre-tRNA processing. The Aq880_Δ181-191_ still forms tetramers and a low number of hexamers but is only 10 % active (**Fig. 2B, C**). In contrast, Aq880_Δ184-191_ also assembles into octamers, decamers and a small subset of even dodecamers but retains over 90 % activity (**Fig. 2B, C, Tab. 1**). We thus consider it likely that higher oligomers form a more rigid scaffold, while a tetramer alone is too flexible for efficient binding and cleavage of precursor tRNAs. HARPs thus represent an impressive example of how a more complex biological task can be accomplished by a small protein that assembles into large homo-oligomeric ultrastructure, and where different monomers contribute distinct partial functions.

**Figure 4.**
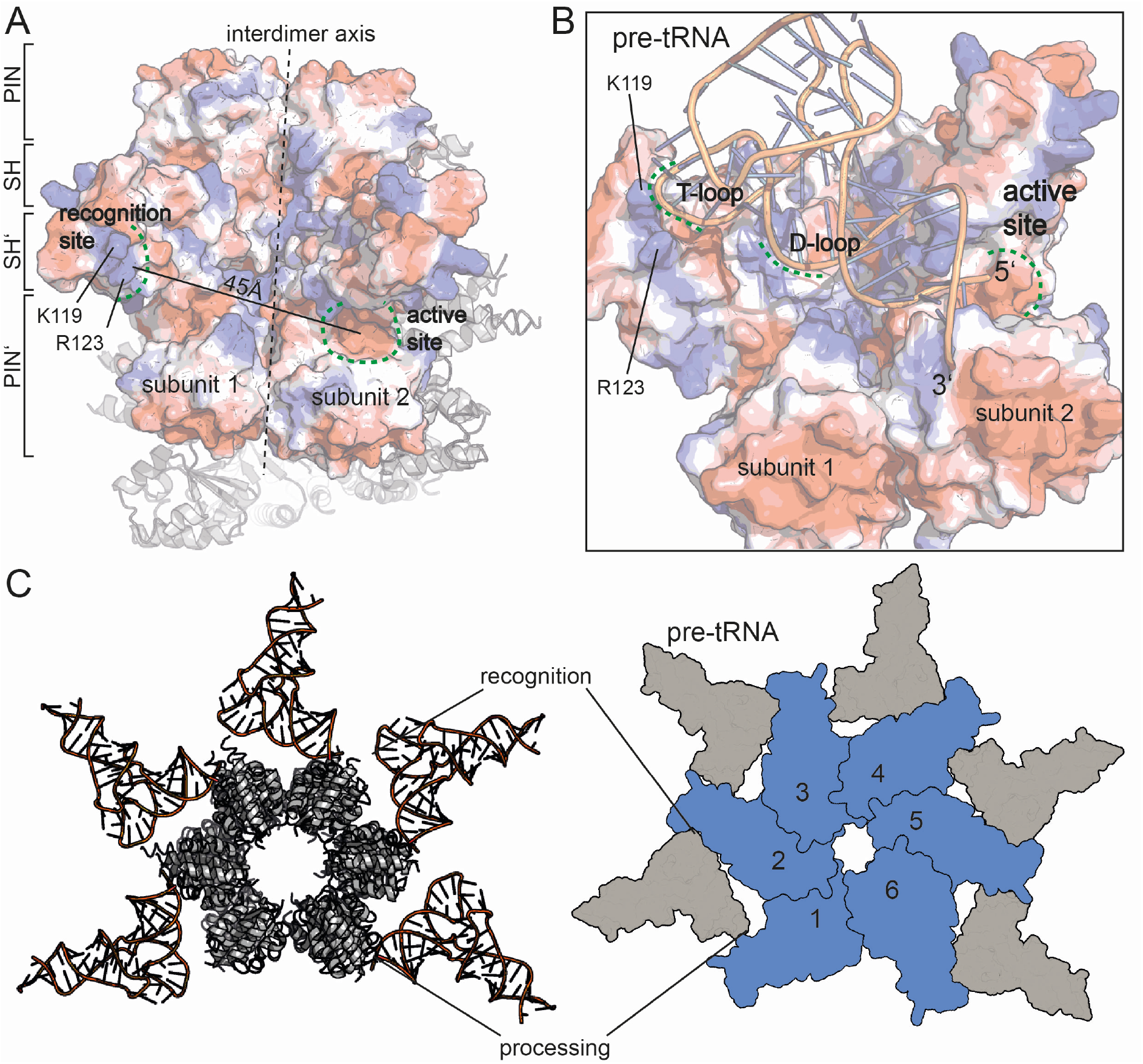
Model for tRNA recognition and processing by HARPs. **A.** Surface of two adjacent Aq880 dimers colored according to the calculated electrostatic potential. The proposed recognition site for the tRNA elbow and the active site are indicated by curved dashed green lines; the almost vertical straight and dashed line marks the interdimer axis. The distance between the two regions is approximately 45 Å. The remaining subunits of the dodecamer are shown as cartoon in the background. **B.** Closeup of the Aq880 surface colored according to the calculated electrostatic potential. The tRNA^Phe^, taken from the PRORP PPR domain co-structure (PDB: 6LVR), was docked onto our structure. The model predicts how the tRNA is coordinated via positive residues within the SH domain to position the 5’-end in close proximity to the active site of the neighboring subunit. **C.** Left: model of the Hhal2243 homo-dodecamer with docked tRNAs. Right: Sketch of tRNA recognition and cleavage by HARPs.

Taken together, we here present the molecular framework for pre-tRNA processing by HARPs and show that this enzyme system, although it evolved independently of PRORPs, shares conserved features with eukaryotic PRORPs concerning pre-tRNA recognition and nuclease activity.

## Material & Methods

### Molecular Cloning and plasmid generation

For Primer and plasmid constructions, see the Supplementary Information (Table S2).

### Expression and purification of recombinant proteins

#### Preparation of Hhal2243

For recombinant overexpression of the HARP from *Halorhodospira halophila* SL1 (Hhal2243) with N-terminal His tag, a pET28(+) Hhal2243nHis plasmid was introduced into Lemo21 (DE3) competent *E. coli* cells as well as into the Rosetta (DE3) strain. For protein expression in the Lemo21 strain, an LB broth culture supplemented with 50 μg/mL kanamycin and 34 μg/mL chloramphenicol was incubated for 6 h at 37°C. A main culture was inoculated 1:50 with this pre-culture supplemented with 500 μM rhamnose. After 2 h at 37°C, overexpression of the Hhal2243nHis protein was induced upon addition of IPTG (0.4 mM) and cells were incubated for another 16 h. Protein expression in the Rosetta strain was done as described below for Aq880 and mutants thereof. Cells obtained in both expression systems were harvested by centrifugation at 2,000 x g for 30 min at 4°C and combined. The combined cell pellets were resuspended in NPI-20 buffer (50 mM NaH_2_PO_4_/NaOH, pH 8.0, 300 mM NaCl, 20 mM imidazole) and disrupted by sonification (output control: 50%, duty cycle: 50%, output: 20%) in 3 cycles for 2 min on ice. After centrifugation (4°C, 1 h, 10,500 g) the lysate was filtered and loaded onto a 1 mL HisTrap column. Elution was performed with NPI-500 buffer (as NPI-20, but containing 500 mM imidazole) applying a linear gradient over 30 column volumes (Fig. S1A), and HARP-containing fractions were dialyzed against thrombin cleavage buffer (10 mM Tris-HCl, pH 8.0, 100 mM KCl, 0.1 mM EDTA, 2 mM CaCl2, 10% (v/v) glycerol). The N-terminal His6-tag was then removed by digestion with thrombin (≈ 1 U/mg; GE Healthcare) at 4°C overnight. For removal of thrombin and further purification, the protein fractions were subjected to MonoQ anion exchange chromatography (Fig. S1A). The tag-free HARP was eluted with thrombin cleavage buffer containing 1 M KCl and dialyzed against “crystallization buffer” (10 mM Tris-HCl pH 8.0, 100 mM KCl, 0.1 mM EDTA). Protein purity was analyzed by 15% SDS-PAGE (Fig. S1B), and successful removal of the His6-tag by thrombin digestion was verified using an α-6His-HRP antibody (Fig. S1C). The protein was frozen in liquid nitrogen and stored at −80°C. All protein concentrations were determined via Bradford assay. All protein preparations were nucleic acid-free based on absorption ratios > 1 for 280/260 nm.

#### Preparation of Aq880 and mutants

Aq880 with C-terminal His6 tag (Aq880cHis) and mutants thereof were overexpressed in Rosetta (DE3) cells in LB autoinduction medium (0.2% (w/v) lactose, 0.05 % (w/v) glucose) supplemented with 50 μg/mL kanamycin and 34 μg/mL chloramphenicol. Cells grown at 37 °C for 18 to 22 h were harvested by centrifugation at 2,000 x g for 30 min at 4 °C. After resuspension in NPI-20 buffer (see above), cells were lysed via sonification (output control: 50%, duty cycle: 50%, output: 20%) in 5 cycles (each 2 min) and with cooling on ice between the cycles. After centrifugation (10,500 x g for 4 °C, 1 h), the lysate was filtered and loaded onto a 1 mL HisTrap column. Elution was performed with NPI-500 buffer applying a step gradient in 20% steps over 20 column volumes. Protein purity was analyzed using 12% stain-free TGX gels detected via the ChemiDoc MP Imaging System (BioRad). The protein was dialyzed against “storage buffer” (10 mM Tris-HCl pH 8.0, 100 mM KCl, 10 mM MgCl_2_, 0.1 mM EDTA, 3 mM DTT immediately added before use, 50% (v/v) glycerol) and stored at −20 °C. For analyzing the protein’s oligomerization state, analytical size exclusion chromatography was performed. Beforehand, Aq880cHis was purified over a MonoQ column and eluting protein fractions were dialyzed against “crystallization buffer” (10 mM Tris-HCl pH 8.0, 100 mM KCl, 0.1 mM EDTA). Then 250 μL protein (0.2 - 2.3 mg) were loaded onto the Superose 6 10/300 GL column. To obtain a calibration curve for molecular mass estimation, protein standards (Merck Sigma-Aldrich) specified in the Supplementary Material were separated on the same column.

#### Cryo-EM grid preparation and data collection

To prepare cryo-EM grids, 3 μL of Hhal2243 at 100 μM concentration were applied to CF 1.2/1.3 grids (Protochips) that were glow-discharged 20 s immediately before use. The sample was incubated 30 s at 100% humidity and 10°C before blotting for 11 s with blotforce −2 and then plunge-frozen into a liquid ethane cooled by liquid nitrogen using a Vitrobot Mark IV (FEI). Data were acquired on a Titan Krios electron microscope (Thermo Fisher Scientific, FEI) operated at 300 kV, equipped with a K3 direct electron detector (Gatan). Movies were recorded in counting mode at a pixel size of 0.833 Å per pixel using a cumulative dose of 40 e-/Å2 and 40 frames. Data acquisition was performed using EPU 2 with two exposures per hole with a target defocus range of 1.5 to 2.4 μm.

#### Cryo-EM data processing

The Hhal2243 dataset was processed in CryoSparc v3.1 (Punjani et al. 2017). Dose-fractionated movies were gain-normalized, aligned, and dose-weighted using Patch Motion correction. The contrast transfer function (CTF) was determined using CTFfind4 (Rohou and Grigorieff 2015). A total of 52,710 particles was picked using the blob picking algorithm and used to train a model that was subsequently used to pick the entire dataset using TOPAZ (Bepler et al. 2019). A total of 2,749,587 candidate particles were extracted and cleaned using iterative-rounds of reference-free 2D classification. The 2,665,011 particles after 2D classification were used for ab initio model reconstruction. The particles were further iteratively classified in 3D using heterogenous refinement. The 1,736,597 particles belonging to the best-aligning particles were subsequently subjected to homogenous 3D refinement, yielding 3.37 Å global resolution and a B-factor of −181.8 Å^2^.

#### Model building

The reconstructed density was interpreted using COOT (Emsley and Cowtan 2004); a model was built manually into the electron density of the best resolved molecule and superposed to reconstruct the symmetry mates. Model building was iteratively interrupted by real-space refinements using Phenix (Liebschner et al. 2019). Statistics assessing the quality of the final model were generated using Molprobity (Chen et al. 2010). Images of the calculated density and the built model were prepared using UCSF Chimera (Pettersen et al. 2004), UCSF ChimeraX (Goddard et al. 2018), and PyMOL.

#### Mass photometry (MP)

Mass photometry experiments were performed using a OneMP mass photometer (Refeyn Ltd, Oxford, UK). Data acquisition was performed using AcquireMP (Refeyn Ltd. v2.3). Mass photometry movies were recorded at 1 kHz, with exposure times varying between 0.6 and 0.9 ms, adjusted to maximize camera counts while avoiding saturation. Microscope slides (70 x 26 mm) were cleaned for 5 min in 50% (v/v) isopropanol (HPLC grade in Milli-Q H_2_O) and pure Milli-Q H_2_O, followed by drying with a pressurized air stream. Silicon gaskets to hold the sample drops were cleaned in the same manner and fixed to clean glass slides immediately prior to measurement. The instrument was calibrated using the NativeMark Protein Standard (Thermo Fisher Scientific) immediately prior to measurements. The concentration during measurement of Aq880, Aq880 mutants or Hhal2243 during measurements was typically 100 nM. Each protein was measured in a new gasket well (i.e., each well was used once). To find focus, 18 μL of fresh buffer adjusted to room temperature was pipetted into a well, the focal position was identified and locked using the autofocus function of the instrument. For each acquisition, 2μL of diluted protein was added to the well and thoroughly mixed. The data were analyzed using the DiscoverMP software.

#### Pre-tRNA processing assays

Activity of recombinant HARPs was analyzed essentially as described (Nickel et al. 2017). Processing assays were carried out in buffer F (50 mM Tris-HCl, pH 7.0, 20 mM NaCl, 5 mM DTT added immediately before use) supplemented with 4.5 mM divalent metal ions (usually 4.5 mM MgCl_2_). Cleavage assays were performed with 50 or 500 nM HARP and ~5 nM 5’-^32^P-labeled pre-tRNA^Gly^. Enzyme and substrate were preincubated separately (enzyme: 5 min at 37 °C; substrate 5 min at 55°C/5 min at 37°C). To start the reaction, 4 μL of substrate mix were added to 16 μL enzyme mix. At different time points, 4 μL aliquots were withdrawn, mixed with 2 × denaturing loading buffer (0.02% (w/v) bromophenol blue, 0.02% (w/v) xylene cyanol blue, 2.6 M urea, 66% (v/v) formamide, 2 × TBE) on ice and subjected to electrophoresis on 20% denaturing polyacrylamide gels. 5’-^32^P-labeled pre-tRNA^Gly^ substrate and the cleaved off 5’-leader product were visualized using a Bio-Imaging Analyzer FLA3000-2R (Fujifilm) and quantified with the AIDA software (Raytest). First-order rate constants of cleavage (*k_obs_*) were calculated with Grafit 5.0.13 (Erithacus Software) by nonlinear regression analysis. HARP working solutions, obtained by dilution from stock solutions, were prepared in EDB buffer (10 mM Tris-HCl pH 7.8, 30 mM NaCl, 0.3 mM EDTA, 1 mM DTT added immediately before use) and kept on ice before use; ~ 1 μL enzyme working solution was added to the aforementioned enzyme mix (Σ 16 μL).

## Acknowledgements

We thank the Cryo-EM facility at the MPI for Biophysics for generous support. We are grateful to Jan Schuller for critical discussions on the manuscript and Cryo-EM data processing. G. H. thanks the Max-Planck Society for financial support. Financial support from the Deutsche Forschungsgemeinschaft to R.K.H. (grant DFG HA 1672/19-1) is acknowledged.

## Author contributions

R.K.H. and F.A. conceived of the project, designed the study and wrote the paper. R.F., N.B.W., S. P., P.I.G. and F.A. performed experiments. R.F., R.K.H and F.A. analyzed data. G.B., G. H. and R.K.H. contributed funding and resources. All authors read and commented on the manuscript.

## Data availability

Coordinates and structure factors have been deposited within the protein data bank (PDB) and the electron microscopy data bank (EMDB) under accession codes: 7OG5 and EMD-12878. The authors declare that all other data supporting the findings of this study are available within the article and its supplementary information files.

## References

Ahnert, S. E., J. A. Marsh, H. Hernandez, C. V. Robinson, and S. A. Teichmann. 2015. “Principles of Assembly Reveal a Periodic Table of Protein Complexes.” Science 350 (6266): aaa2245–aaa2245. https://doi.org/10.1126/science.aaa2245.

Bazin, Alexandre, Mickaël V Cherrier, Irina Gutsche, Joanna Timmins, and Laurent Terradot. 2015. “Structure and Primase-Mediated Activation of a Bacterial Dodecameric Replicative Helicase.” Nucleic Acids Research 43 (17): 8564–76. https://doi.org/10.1093/nar/gkv792.

Bepler, Tristan, Andrew Morin, Micah Rapp, Julia Brasch, Lawrence Shapiro, Alex J. Noble, and Bonnie Berger. 2019. “Positive-Unlabeled Convolutional Neural Networks for Particle Picking in Cryo-Electron Micrographs.” Nature Methods 16 (11): 1153–60. https://doi.org/10.1038/s41592-019-0575-8.

Breuert, Sebastian, Thorsten Allers, Gabi Spohn, and Jörg Soppa. 2006. “Regulated Polyploidy in Halophilic Archaea.” Edited by Beth Sullivan. PLoS ONE 1 (1): e92. https://doi.org/10.1371/journal.pone.0000092.

Chen, Vincent B., W. Bryan Arendall, Jeffrey J. Headd, Daniel A. Keedy, Robert M. Immormino, Gary J. Kapral, Laura W. Murray, Jane S. Richardson, and David C. Richardson. 2010. “MolProbity: All-Atom Structure Validation for Macromolecular Crystallography.” Acta Crystallographica Section D: Biological Crystallography 66 (1): 12–21. https://doi.org/10.1107/S0907444909042073.

Daniels, Charles J., Lien B. Lai, Tien Hao Chen, and Venkat Gopalan. 2019. “Both Kinds of RNase P in All Domains of Life: Surprises Galore.” RNA 25 (3): 286–91. https://doi.org/10.1261/rna.068379.118.

Emsley, Paul, and Kevin Cowtan. 2004. “Coot: Model-Building Tools for Molecular Graphics.” Acta Crystallographica. Section D, Biological Crystallography 60 (Pt 12 Pt 1): 2126–32. https://doi.org/10.1107/S0907444904019158.

Gobert, Anthony, Mathieu Bruggeman, and Philippe Giegé. 2019. “Involvement of PIN-like Domain Nucleases in TRNA Processing and Translation Regulation.” IUBMB Life. Blackwell Publishing Ltd. https://doi.org/10.1002/iub.2062.

Gobert, Anthony, Bernard Gutmann, Andreas Taschner, Markus Göringer, Johann Holzmann, Roland K. Hartmann, Walter Rossmanith, and Philippe Giegé. 2010. “A Single Arabidopsis Organellar Protein Has RNase P Activity.” Nature Structural and Molecular Biology 17 (6): 740–44. https://doi.org/10.1038/nsmb.1812.

Goddard, Thomas D., Conrad C. Huang, Elaine C. Meng, Eric F. Pettersen, Gregory S. Couch, John H. Morris, and Thomas E. Ferrin. 2018. “UCSF ChimeraX: Meeting Modern Challenges in Visualization and Analysis.” Protein Science 27 (1): 14–25. https://doi.org/10.1002/pro.3235.

Guerrier-Takada, Cecilia, Katheleen Gardiner, Terry Marsh, Norman Pace, and Sidney Altman. 1983. “The RNA Moiety of Ribonuclease P Is the Catalytic Subunit of the Enzyme.” Cell 35 (3 PART 2): 849–57. https://doi.org/10.1016/0092-8674(83)90117-4.

Holzmann, Johann, Peter Frank, Esther Löffler, Keiryn L. Bennett, Christopher Gerner, and Walter Rossmanith. 2008. “RNase P without RNA: Identification and Functional Reconstitution of the Human Mitochondrial TRNA Processing Enzyme.” Cell 135 (3): 462–74. https://doi.org/10.1016/j.cell.2008.09.013.

Howard, Michael J., Wan Hsin Lim, Carol A. Fierke, and Markos Koutmos. 2012. “Mitochondrial Ribonuclease P Structure Provides Insight into the Evolution of Catalytic Strategies for Precursor-TRNA 5’ Processing.” Proceedings of the National Academy of Sciences of the United States of America 109 (40): 16149–54. https://doi.org/10.1073/pnas.1209062109.

Jarrous, Nayef, and Venkat Gopalan. 2010. “Archaeal/Eukaryal RNase P: Subunits, Functions and RNA Diversification.” Nucleic Acids Research 38 (22): 7885–94. https://doi.org/10.1093/nar/gkq701.

Klemm, Bradley P., Nancy Wu, Yu Chen, Xin Liu, Kipchumba J. Kaitany, Michael J. Howard, and Carol A. Fierke. 2016. “The Diversity of Ribonuclease P: Protein and RNA Catalysts with Analogous Biological Functions.” Biomolecules. MDPI AG. https://doi.org/10.3390/biom6020027.

Laniado, Joshua, and Todd O. Yeates. 2020. “A Complete Rule Set for Designing Symmetry Combination Materials from Protein Molecules.” Proceedings of the National Academy of Sciences of the United States of America 117 (50): 31817–23. https://doi.org/10.1073/pnas.2015183117.

Lechner, Marcus, Walter Rossmanith, Roland K. Hartmann, Clemens Thölken, Bernard Gutmann, Philippe Giegé, and Anthony Gobert. 2015. “Distribution of Ribonucleoprotein and Protein-Only RNase P in Eukarya.” Molecular Biology and Evolution 32 (12): 3186–93. https://doi.org/10.1093/molbev/msv187.

Liebschner, Dorothee, Pavel V. Afonine, Matthew L. Baker, Gábor Bunkoczi, Vincent B. Chen, Tristan I. Croll, Bradley Hintze, et al. 2019. “Macromolecular Structure Determination Using X-Rays, Neutrons and Electrons: Recent Developments in Phenix.” Acta Crystallographica Section D: Structural Biology 75 (Pt 10): 861–77. https://doi.org/10.1107/S2059798319011471.

Marszalkowski, Michal, Dagmar K. Willkomm, and Roland K. Hartmann. 2008. “5’-End Maturation of TRNA in Aquifex Aeolicus.” Biological Chemistry 389 (4): 395–403. https://doi.org/10.1515/BC.2008.042.

Masuda, Hisako, and Masayori Inouye. 2017. “Toxins of Prokaryotic Toxin-Antitoxin Systems with Sequence-Specific Endoribonuclease Activity.” Toxins 9 (4): 140. https://doi.org/10.3390/toxins9040140.

Matelska, Dorota, Kamil Steczkiewicz, and Krzysztof Ginalski. 2017. “Comprehensive Classification of the PIN Domain-like Superfamily.” Nucleic Acids Research. Oxford University Press. https://doi.org/10.1093/nar/gkx494.

Nickel, Astrid I., Nadine B. Wäber, Markus Gößringer, Marcus Lechner, Uwe Linne, Ursula Toth, Walter Rossmanith, and Roland K. Hartmann. 2017. “Minimal and RNA-Free RNase P in Aquifex Aeolicus.” Proceedings of the National Academy of Sciences of the United States of America 114 (42): 11121–26. https://doi.org/10.1073/pnas.1707862114.

Pettersen, Eric F., Thomas D. Goddard, Conrad C. Huang, Gregory S. Couch, Daniel M. Greenblatt, Elaine C. Meng, and Thomas E. Ferrin. 2004. “UCSF Chimera? A Visualization System for Exploratory Research and Analysis.” Journal of Computational Chemistry 25 (13): 1605–12. https://doi.org/10.1002/jcc.20084.

Punjani, Ali, John L. Rubinstein, David J. Fleet, and Marcus A. Brubaker. 2017. “CryoSPARC: Algorithms for Rapid Unsupervised Cryo-EM Structure Determination.” Nature Methods 14 (3): 290–96. https://doi.org/10.1038/nmeth.4169.

Rohou, Alexis, and Nikolaus Grigorieff. 2015. “CTFFIND4: Fast and Accurate Defocus Estimation from Electron Micrographs.” Journal of Structural Biology 192 (2): 216–21. https://doi.org/10.1016/j.jsb.2015.08.008.

Rooyen, Jason M. Van, Valerie R. Abratt, Hassan Belrhali, and Trevor Sewell. 2011. “Crystal Structure of Type III Glutamine Synthetase: Surprising Reversal of the Inter-Ring Interface.” Structure 19 (4): 471–83. https://doi.org/10.1016/j.str.2011.02.001.

Roy, Siddhartha, Ramachandran Saraswathi, Dipankar Chatterji, and M. Vijayan. 2008. “Structural Studies on the Second Mycobacterium Smegmatis Dps: Invariant and Variable Features of Structure, Assembly and Function.” Journal of Molecular Biology 375 (4): 948–59. https://doi.org/10.1016/j.jmb.2007.10.023.

Schencking, Isabell, Walter Rossmanith, and Roland K. Hartmann. 2020. “Diversity and Evolution of RNase P.” In Evolutionary Biology—A Transdisciplinary Approach, 255–99. Springer International Publishing. https://doi.org/10.1007/978-3-030-57246-4_11.

Schwarz, Thandi S., Nadine B. Wäber, Rebecca Feyh, Katrin Weidenbach, Ruth A. Schmitz, Anita Marchfelder, and Roland K. Hartmann. 2019. “Homologs of Aquifex Aeolicus Protein-Only RNase P Are Not the Major RNase P Activities in the Archaea Haloferax Volcanii and Methanosarcina Mazei.” IUBMB Life 71 (8): 1109–16. https://doi.org/10.1002/iub.2122.

Soltermann, Fabian, Eric D.B. Foley, Veronica Pagnoni, Martin Galpin, Justin L.P. Benesch, Philipp Kukura, and Weston B. Struwe. 2020. “Quantifying Protein–Protein Interactions by Molecular Counting with Mass Photometry.” Angewandte Chemie - International Edition 59 (27): 10774–79. https://doi.org/10.1002/anie.202001578.

Sonn-Segev, Adar, Katarina Belacic, Tatyana Bodrug, Gavin Young, Ryan T VanderLinden, Brenda A Schulman, Johannes Schimpf, et al. 2020. “Quantifying the Heterogeneity of Macromolecular Machines by Mass Photometry.” Nature Communications 11 (1): 1772. https://doi.org/10.1038/s41467-020-15642-w.

Stachler, Aris Edda, and Anita Marchfelder. 2016. “Gene Repression in Haloarchaea Using the CRISPR (Clustered Regularly Interspaced Short Palindromic Repeats)-Cas I-B System.” Journal of Biological Chemistry 291 (29): 15226–42. https://doi.org/10.1074/jbc.M116.724062.

Teramoto, Takamasa, Kipchumba J. Kaitany, Yoshimitsu Kakuta, Makoto Kimura, Carol A. Fierke, and Traci M. Tanaka Hall. 2020. “Pentatricopeptide Repeats of Protein-Only RNase P Use a Distinct Mode to Recognize Conserved Bases and Structural Elements of Pre-TRNA.” Nucleic Acids Research 48 (21): 11815–26. https://doi.org/10.1093/nar/gkaa627.

Yan, Junjie, Yinying Yao, Sixing Hong, Yan Yang, Cuicui Shen, Qunxia Zhang, Delin Zhang, Tingting Zou, and Ping Yin. 2019. “Delineation of Pentatricopeptide Repeat Codes for Target RNA Prediction.” Nucleic Acids Research 47 (7): 3728–38. https://doi.org/10.1093/nar/gkz075.

