## Supplementary Material for "Structure and mechanistic features of the prokaryotic minimal RNase P"

**Supplementary Table 1. Cryo-EM data collection, refinement and validation statistics.**

| | Hhal2243 (EMD-12878) (PDB: 7OG5) |
|---|---|
| **Data collection and processing** | |
| Magnification | 105,000 |
| Voltage (kV) | 300 |
| Electron exposure (e-/Å$^2$) | 40 |
| Defocus range (μm) | 1.5 – 2.4 |
| Pixel size (Å) | 0.833 |
| Symmetry imposed | C1 |
| Initial particle images (no.) | 2,749,587 |
| Final particle images (no.) | 1,736,597 |
| Map Resolution (Å) | 3.37 |
| FSC threshold | 0.143 |
| Map resolution range | 3.37 – 5 Å |
| **Refinement** | |
| Map sharpening *B* factor (Å$^2$) | 181 |
| Model composition | |
| Non-hydrogen atoms | 15444 |
| Protein residues | 1892 |
| Ligands | 0 |
| *B* factors (Å$^2$) | |
| Protein | 106.70 |
| Ligand | 0 |
| R.m.s. deviations | |
| Bond Lengths (Å) | 0.01 |
| Bond angles (°) | 0.933 |
| Validation | |
| MolProbity score | 2.33 |
| Clashscore | 19.16 |
| Poor rotamers (%) | 0 |
| Ramachandran plot | |
| Favored (%) | 89.91 |
| Allowed (%) | 10.09 |
| Outliers (%) | 0.00 |

*Molecular Cloning and plasmid generation.* In order to overexpress the Aq880cHis protein a pET-28a(+)_Aq880cHis plasmid was constructed. For this purpose, the genomic DNA of *A. aeolicus* strain was isolated and used as template for amplification of the aq_880 gene with the primer pair listed in table 2 as described by (1). The PCR product was inserted into the pET-28a(+) vector via *Xho*I and *Nco*I restriction sites. The tag-free Aq880 construct was cloned by using the pET-28a(+)_Aq880cHis plasmid as template and the appropriate primer pair to remove the His6-tag. For all the Aq880cHis variants site-directed mutagenesis was done to gain either the C-terminal truncated proteins or the arginine and lysine-to-alanine mutants. The PCR was performed using the Platinum SuperFi PCR Master Mix (Invitrogen) and the primers listed in table 2. For the pET-28a(+)_Hhal2243nHis construct isolated chromosomal DNA of *H. halophila* was used as template and the PCR amplified product (primers listed below) was inserted via the restriction sites *Nhe*I and *Bpu1102*I into the pET-28a(+) vector (2).

**Supplementary Table 2**: **Primers & Plasmids used in this study**

| constructs in pET-28a(+) | sequence (5'→ 3') |
|---|---|
| aq880 | AAG CCA TGG ATG TGT TCG TTC TCG ACA C &<br>CTC TCG AGA AAC CTG TGT CTT ACC AAG C |
| aq880_Δ184-191 | TTT CTC GAG CAC CAC CAC CAC CAC C &<br>AATGTTTTTGAAATTCTTAGGGTCTATG |
| aq880_Δ181-191 | TTT CTC GAG CAC CAC CAC CAC CAC C &<br>GAA ATT CTT AGG GTC TAT GAG TTT TAT ACC |
| aq880_Δ179-191 | TTT CTC GAG CAC CAC CAC CAC CAC C &<br>CTT AGG GTC TAT GAG TTT TAT ACC TAT C |
| aq880_Δ177-191 | TTT CTC GAG CAC CAC CAC CAC CAC C &<br>GTC TAT GAG TTT TAT ACC TAT CTT GTC CG |
| aq880_Δ172-191 | TTT CTC GAG CAC CAC CAC CAC CAC C &<br>TAT CTT GTC CGC CCA TGT TCT GAG GCC |
| aq880_K119A/R123A/<br>R125A/K127A/R129A | GCG CAG CTA TTA ATG CCC CGA CGT CTT CAC &<br>TCG CGG AGG CGT ACG CGG AAG CCC TCA GG |
| aq880_R125A/R129A | GCC TAG CTA TTA ATT TCC CGA CGT CTT CAC &<br>TCG CGG AGA AGT ACG CGG AAG CCC TCA GG |
| aq880_R125A | GCC TAG CTA TTA ATT TCC CGA CGT CTT CAC &<br>TCG CGG AGA AGT ACA GGG AAG CCC TCA GG |
| aq880_R129A | GCC TAG CTA TTA ATT TCC CGA CGT CTT CAC &<br>TCA GGG AGA AGT ACG CGG AAG CC |
| hhal2243 | TTT GCT AGC CGC CGA TTC GTG CTC G &<br>TTT GCT CAG CCT ACC CGG CGG GCT G |

**Supplementary Table 3**: Dynamic mass distribution determined by mass photometry

| Sample | Peak | Molecular weight (kDa) | Amount (%) |
|---|---|---|---|
| **Hhal2243 wt** | 1 | 295 | 88 |
| **Aq880 wt** | 1 | 42 | 7 |
| | 2 | 78 | 13 |
| | 3 | 129 | 9 |
| | 4 | 182 | 13 |
| | 5 | 234 | 11 |
| | 6 | 277 | 47 |
| **Aq880_Δ184-191** | 1 | 42 | 32 |
| | 2 | 83 | 50 |
| | 3 | 133 | 12 |
| | 4 | 183 | 4 |
| | 5 | 235 | 2 |
| **Aq880_Δ181-191** | 1 | 42 | 55 |
| | 2 | 81 | 41 |

| | | | |
|---|---|---|---|
| | 3 | 134 | 4 |
| **Aq880_Δ179-191** | 1 | 44 | 90 |
| **Aq880_Δ177-191** | 1 | 52 | 92 |

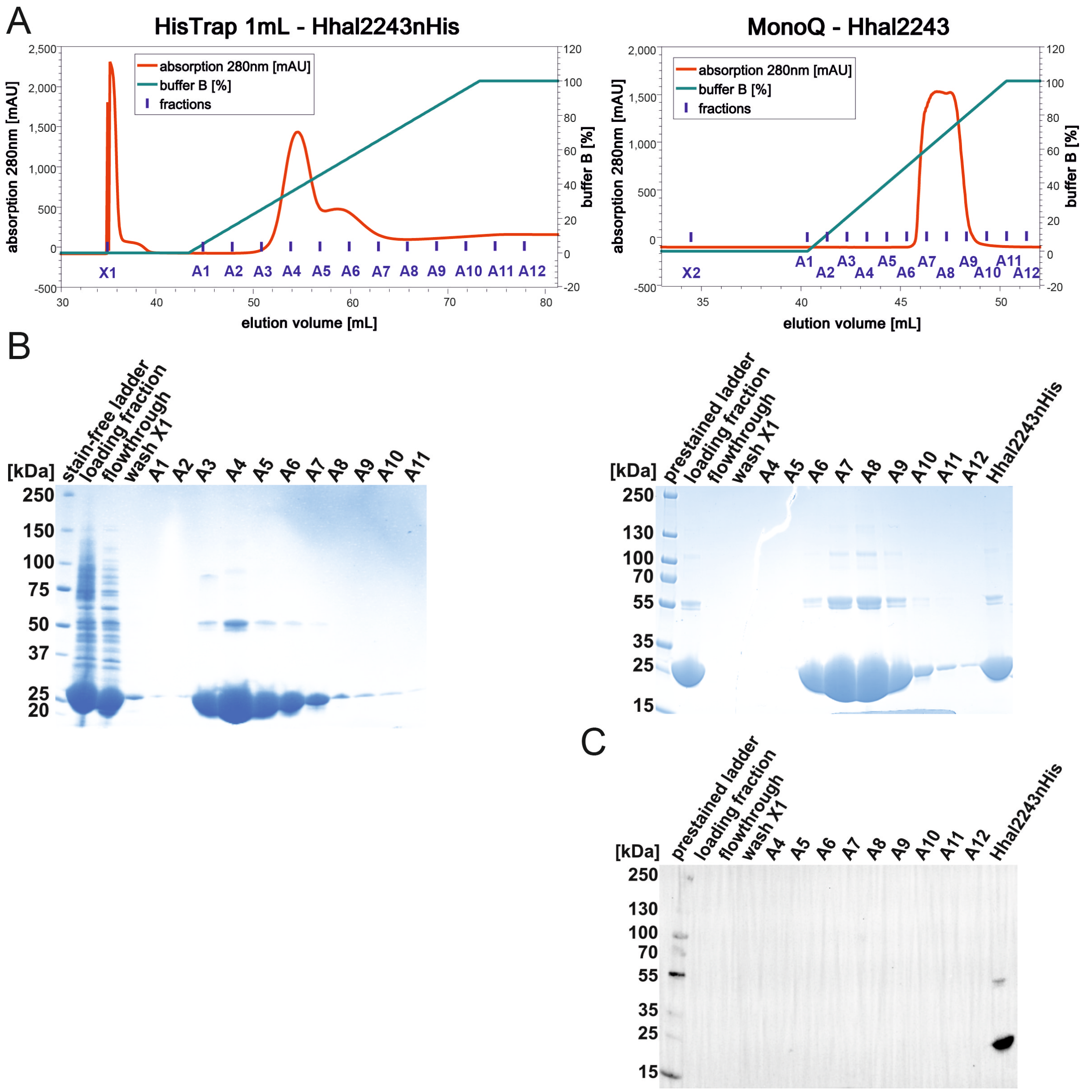

**Supplementary Figure 1. Purification of Hhal2243. A.** Chromatograms of the Ni-ion-affinity chromatography (HisTrap 1mL column, left) and ion-exchange chromatography (MonoQ column, right). **B.** SDS-gels of fractions from the HisTrap (left) and MonoQ column (right). **C.** Western Blot analysis of fractions from the MonoQ column to verify successful removal of the His6-tag by thrombin digestion using an α-6His-HRP antibody (Thermo Fisher Scientific Invitrogen) diluted 1:5000 in 1 % Casein Blocker (Thermo Fisher Scientific).

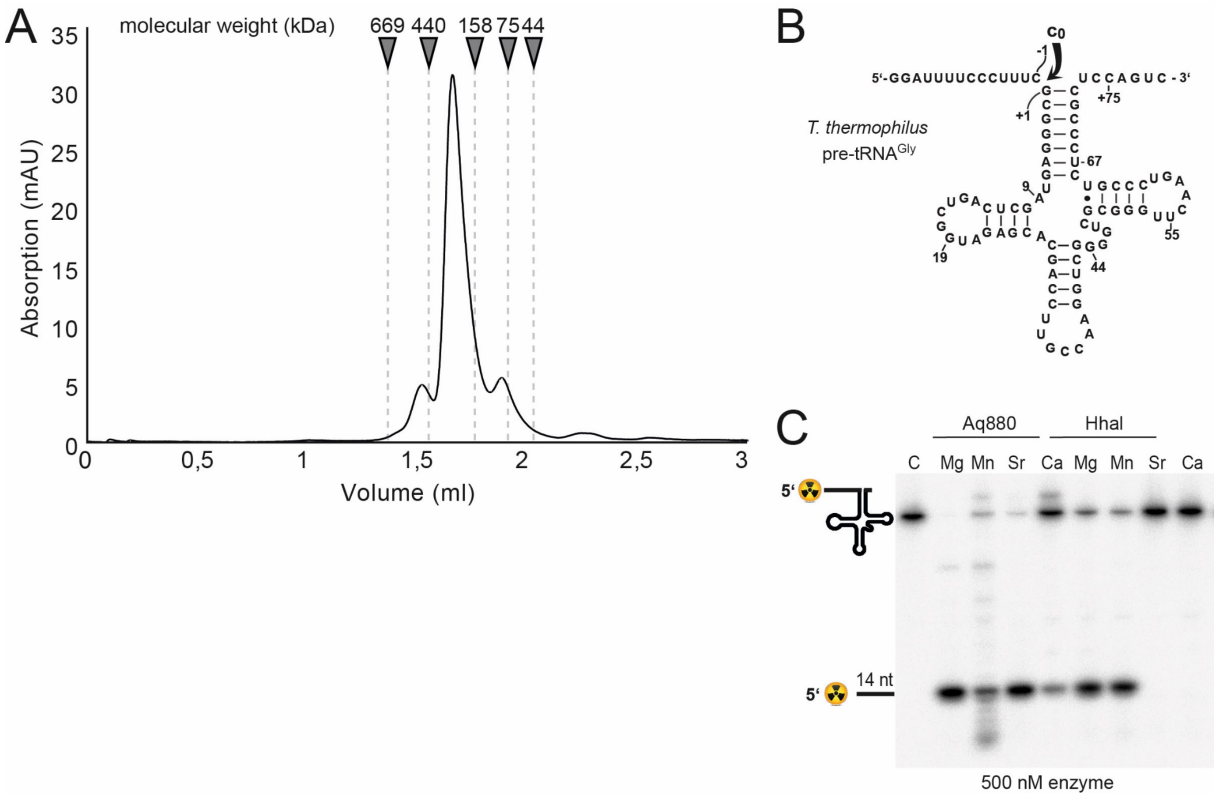

**Supplementary Figure 2.** **A.** Chromatographic analysis of Hhal2243. The sample was analyzed on a Superose 6 3.2/300 column using an ÄKTA PURE (GE Healthcare) FPLC system operated at room temperature. **B.** Secondary structure of precursor tRNA$^{Gly}$ (pre-tRNA$^{Gly}$) of the thermophilic bacterium *Thermus thermophilus* carrying a 5'-leader of 14 nucleotides; the curved arrow indicates the canonical ($c_0$) RNase P cleavage site **C.** RNase P processing of 5'-$^{32}$P-labeled *T. thermophilus* pre-tRNA$^{Gly}$ by recombinant HARPs from the hyperthermophilic bacterium *A. aeolicus* (Aq880) and the γ-proteobacterium *Halorhodospira halophila* (Hhal2243) in the presence of either 4.5 mM $Mg^{2+}$, $Mn^{2+}$, $Sr^{2+}$ or $Ca^{2+}$ in buffer F for 2 h at 37 °C. Enzyme and substrate concentrations were 500 nM and ~5 nM, respectively. Samples were separated by 20 % denaturing PAGE and analyzed by phosphorimaging (for details, see Materials and Methods of the main text). C, control, substrate alone incubated for 2 h at 37 °C in enzyme dilution buffer (30 mM Tris-HCl pH 7.8, 30 mM NaCl, 0.3 mM EDTA, 1 mM DTT).

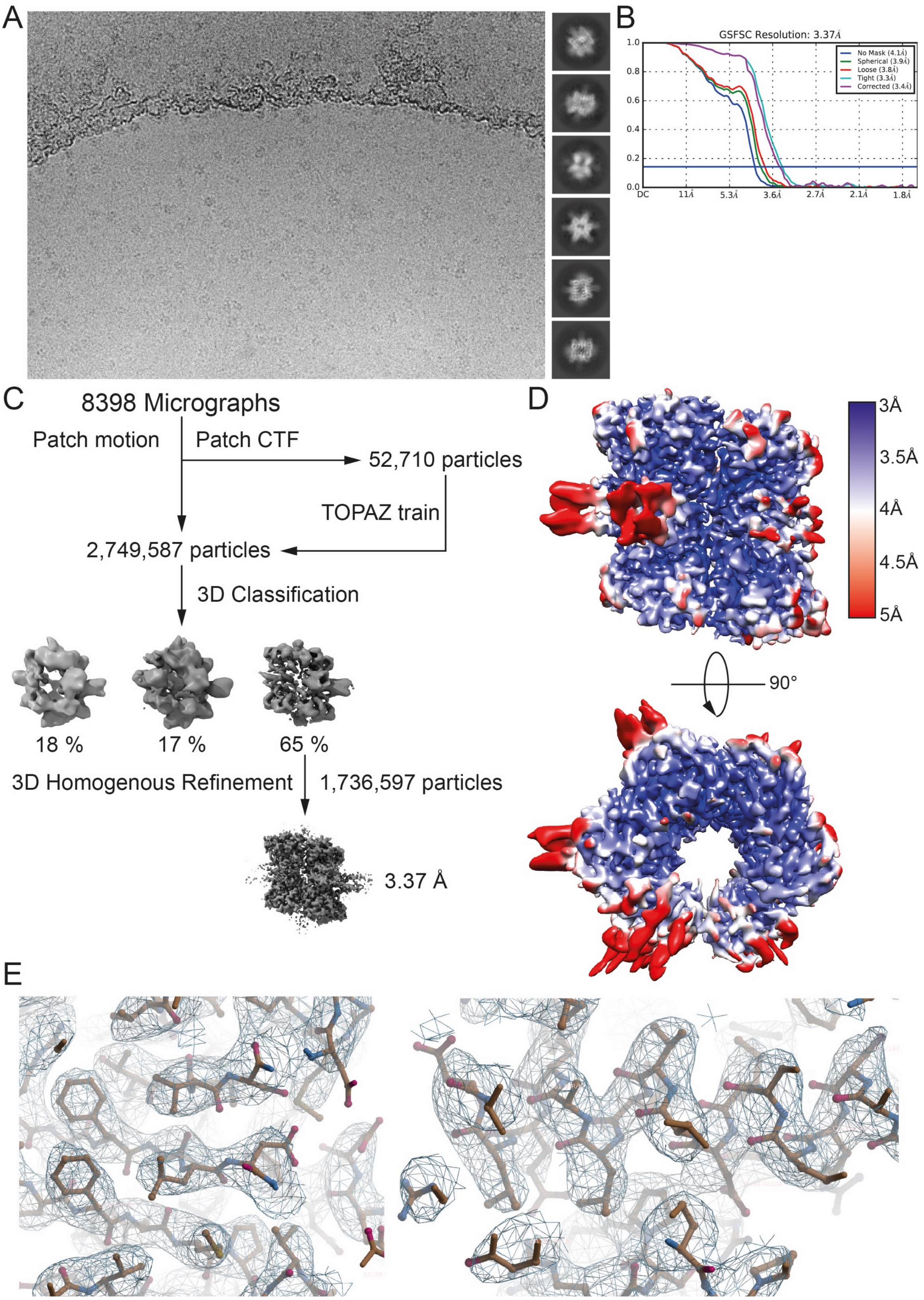

**Supplementary Figure 3. Cryo-EM data collection and analysis. A.** Representative cryo-EM micrograph collected with a Thermo Fisher Scientific Titan Krios microscope, operated at 300 kV and equipped with a K3 direct electron detector. Representative reference-free 2D class averages are shown on the right side of the micrograph. **B.** FSC analysis of the dataset with the blue line representing the estimated global FSC of 3.4 Å. **C.** The 2,749,587 particles from the dataset were 3D classified into three classes using a reference *ab initio* model generated in Cryosparc v3.1. The class consisting of the best aligning particles was selected for 3D homogenous refinement in Cryosparc resulting in a final map at 3.37 Å resolution. **D.** Local resolution estimation of the final map colored from blue (3 Å) to red (5 Å). **E.** Representative images of the Hhal2243 model and surrounding electron density. Maps are displayed as mesh and contoured at 3$\sigma$.

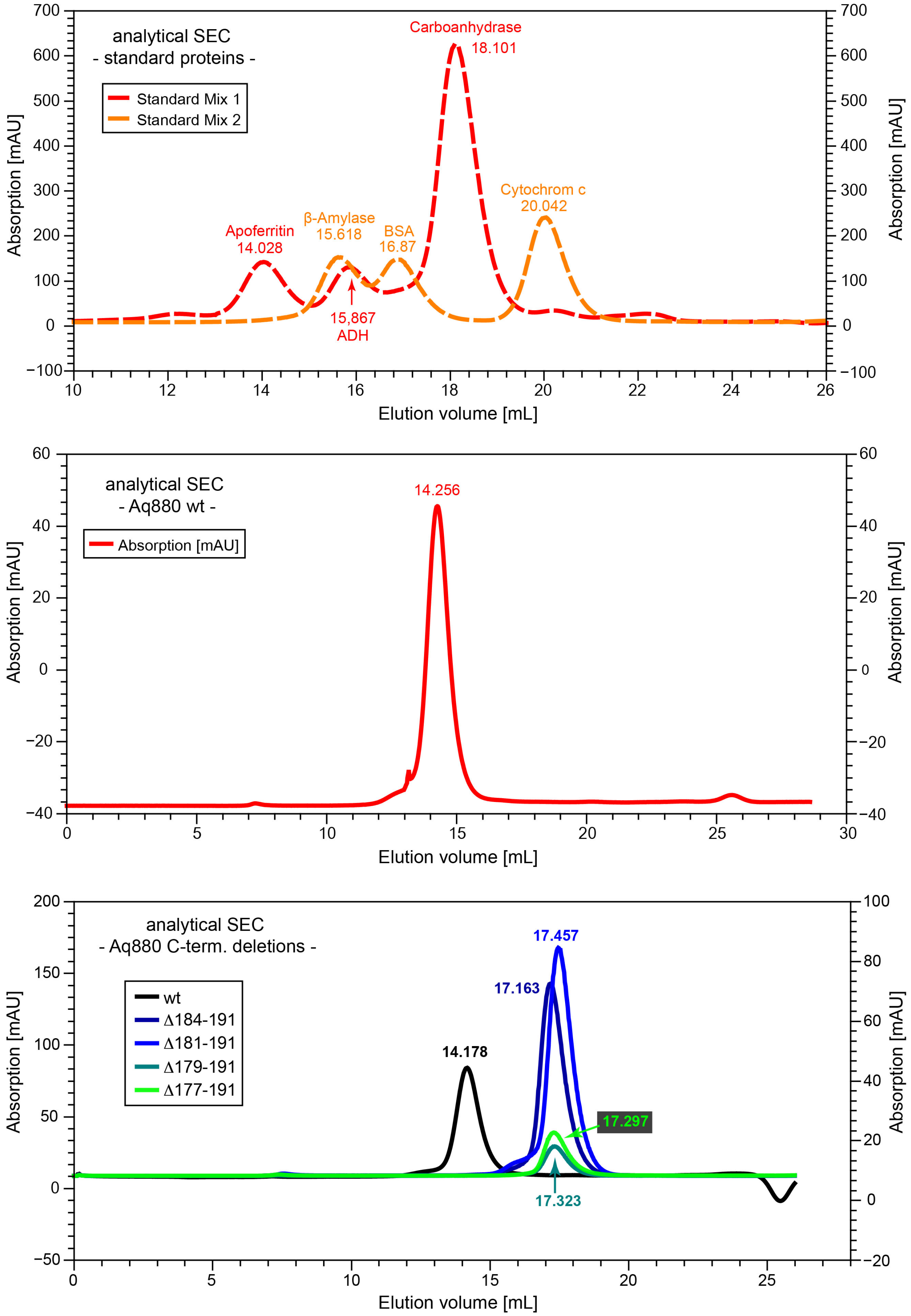

**Supplementary Figure 4: Chromatographic analysis of Aq880 WT and C-terminal deletion mutants.** Samples were analyzed on a Superose 6 10/300 GL column using an ÄKTA Purifier (GE Healthcare) FPLC system operated at room temperature. The running buffer contained 20 mM Tris-HCl pH 8.0, 100 mM KCl, 0.1 mM EDTA and 3 mM DTT. The flow rate was 0.5 mL/min. **A**. Overlay of two column runs with three marker proteins each, termed Standard Mix 1 and 2 (Gel Filtration Markers Kit for Protein Molecular Weights 12-200 kDa and Apoferritin, Merck Sigma-Aldrich). Identities of individual proteins and elution volumes of the respective peak fractions are indicated; ADH, alcohol dehydrogenase; BSA, bovine serum albumin. **B**. Elution profile of Aq880 wt. **C**. Elution profiles of C-terminal deletion variants of Aq880 lacking the indicated amino acids of the native protein. The elution profiles in panel A suggests an approximate mass of 430 kDa for Aq880 wt. The elution volumes of the C-terminal deletions mutants suggest molecular masses between ~55 and 68 kDa.

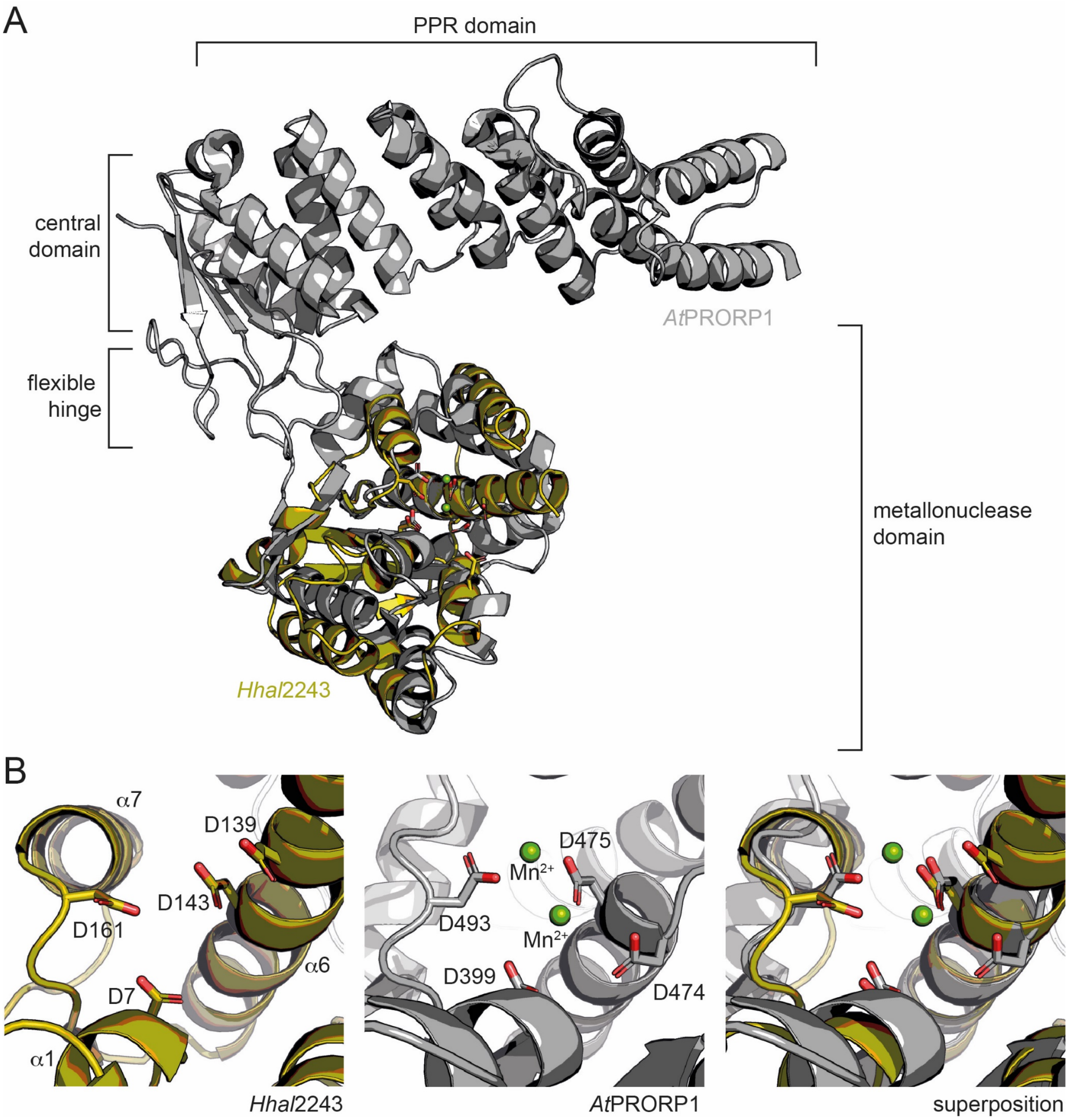

**Supplementary Figure 5. Comparison between HARP and PRORP systems. A.** Superposition of *At*PRORP (PDB: 4G24) and Hhal2243 shows that the metallonuclease domain superposes well, while the PPR domain is absent in HARPs. **B.** Orientation of active site residues is conserved among PRORPs and HARPs; the active site residues are positioned similarly in both systems. AtPRORP is colored in grey and Hhal2243 in olive.

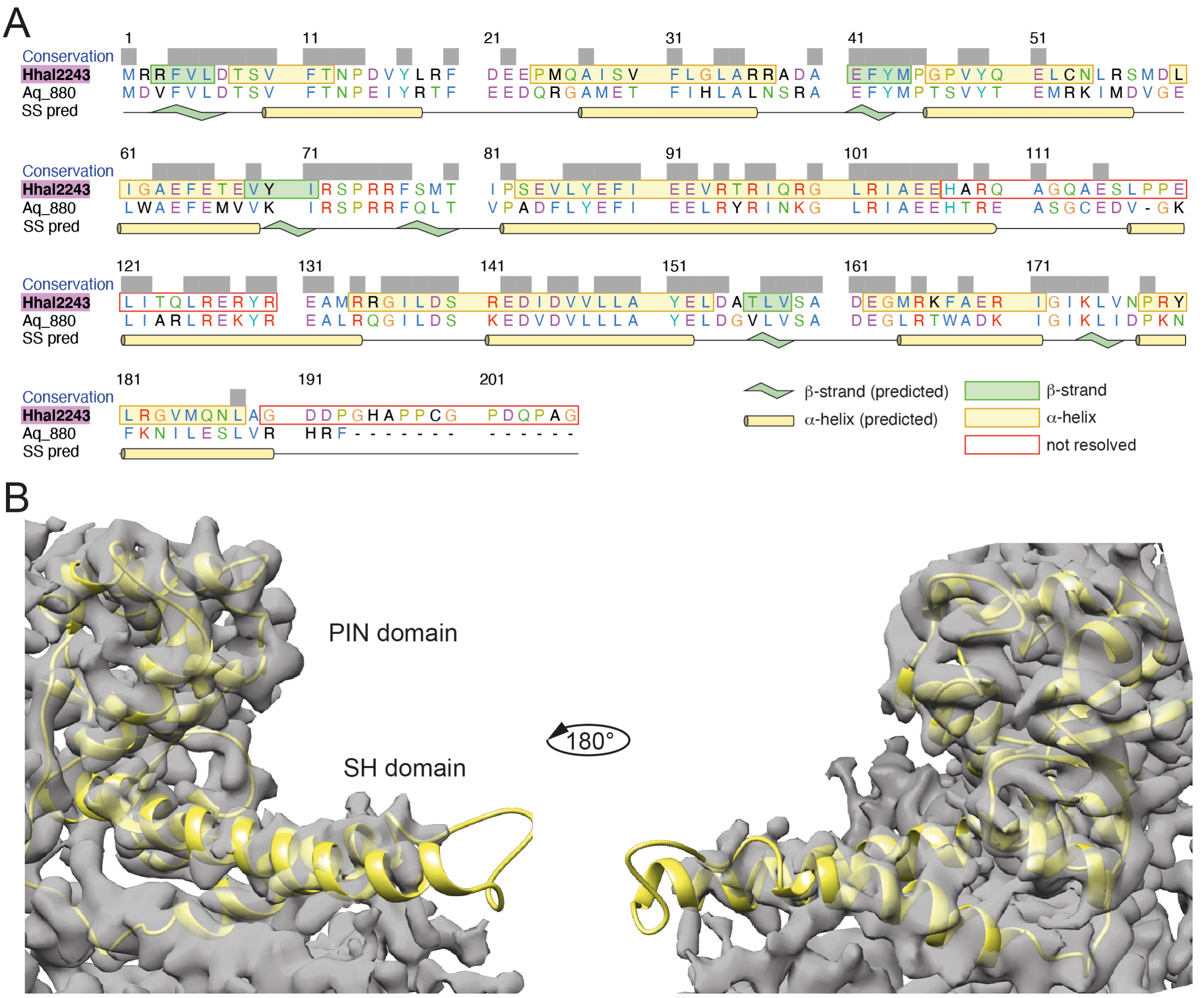

**Supplementary Figure 6. A.** Sequence alignment of Hhal2243 and Aq880. Secondary structure elements are drawn as observed in the Cryo-EM structure of Hhal2243. The secondary structure prediction has been generated using PSIPRED (3) and is shown below the sequence alignment. **B.** Electron density map contoured at level 0.12 with the docked Aq880 model shown in yellow.

**Supplementary References**

1. Nickel AI, Wäber NB, Gößringer M, Lechner M, Linne U, Toth U, Rossmanith W, Hartmann RK (2017) Minimal and RNA-free RNase P in Aquifex aeolicus. *Proc Natl Acad Sci U S A* 114(42):11121–11126.
2. Schwarz TS, Wäber NB, Feyh R, Weidenbach K, Schmitz RA, Marchfelder A, Hartmann RK (2019) Homologs of aquifex aeolicus protein-only RNase P are not the major RNase P activities in the archaea haloferax volcanii and methanosarcina mazei. *IUBMB Life* 71(8):1109–1116.
3. Buchan DWA, Jones DT (2019) The PSIPRED Protein Analysis Workbench: 20 years on. *Nucleic Acids Res* 47(W1):W402–W407.
